# Reciprocal differentiation via GABAergic components and ASD-related phenotypes in hES with 1q21.1 CNV

**DOI:** 10.1101/2021.09.13.460033

**Authors:** Yoshiko Nomura, Jun Nomura, Toru Nishikawa, Toru Takumi

## Abstract

Copy number variations (CNVs) in the distal 1q21.1 region, both deletion (1q del) and duplication (1q dup), are associated with autism spectrum disorder, epilepsy and schizophrenia. Besides common phenotypes, 1q del and 1q dup manifest opposite clinical phenotypes—e.g., microcephaly in 1q del and macrocephaly in 1q dup. However, molecular and cellular mechanisms are still elusive. We generate isogenic human ES (hES) cell lines with reciprocal 1q21.1 CNVs using CRISPR/Cas9 system and differentiate them into 2-dimensional (2-D) neurons and 3-D cortical organoids. Our study recapitulates opposite organoid size and shows dosage-dependent differentiation changes i.e., more mature and GABAergic components in 1q del and more proliferative state in 1q dup. In contrast, both CNVs show hyperexcitability and altered expressions of glutamate system as common features. These results demonstrate that 1q21.1 CNVs dramatically affect cell fate in the early neurodevelopmental periods. This is the first isogenic model of hES CNVs and our findings provide new insights into the underlying mechanisms of neurodevelopmental disorders.

## Introduction

Autism spectrum disorder (ASD) is one of the most common neurodevelopmental diseases and is characterized by persistent deficits in social communication and interaction across multiple contexts and restricted, repetitive patterns of behavior, interests, or activities (Lord *et al*, 2018). Despite its high prevalence (0.6-2.6%) (Maenner *et al*, 2020; Lyall *et al*, 2017) and the significant impacts on one’s social life, biological mechanisms for developing ASD are not well understood and no fundamental treatments have been established.

Although the exact cause of ASD is still unknown, genetic and environmental factors and their interactions have been considered in the context of ASD etiology (Lyall *et al*, 2017; De La Torre-Ubieta *et al*, 2016; Huguet *et al*, 2013; Voineagu *et al*, 2011). Genetic analyses have revealed that different recurrent copy number variations (CNVs) located on chromosome loci such as 1q21.1, 3q29, 7q11.23, 15q11-13, 16p11.2 and 22q11.2 are associated with ASD, suggesting their possible implication in the susceptibility and/or pathophysiology of ASD(Takumi & Tamada, 2018; Malhotra & Sebat, 2012; Bucan *et al*, 2009). Since most of the genes mapped to the above-mentioned chromosomal regions are not specifically linked to ASD and CNVs in these loci are also linked to other psychiatric disorders including schizophrenia and bipolar disorder, various ASD-related CNVs, which are presumed to have respective unique structures, could be involved in a variety of ASD symptoms by exerting distinct effects on the control of gene expression and its neuronal and behavioral consequences. Indeed, previous studies using chromosome-engineered mouse models of human ASD-related CNVs on chromosome 1q21.1, 7q11.23, 15q11-13, 15q13.3, 16p11.2, and 17p11.2 have demonstrated that the respective chromosome-modified mice differentially mimicked human ASD-correlated disturbances in social and spontaneous behavior, sensorimotor processing and functioning, motor coordination, sleep or circadian rhythm, brain morphology, and neurochemical or histological conditions (Takumi *et al*, 2020).

As for comorbidities of ASD, epilepsy is one of the most closely associated symptoms with ASD (2.4-26%) and previous reports pointed out that nearly half of all ASD patients developed epilepsy (El Achkar & Spence, 2015). A large clinical study reported that epilepsy could be associated with exacerbation of some behavioral symptoms in ASD patients (Viscidi *et al*, 2014). Alteration of the brain size is also common in patients with ASD; meta-analysis of the brain size measurement data in ASD has indicated that brain growth outside the normal range in ASD patients might emerge in the first years of life from the age of 2 to 5 (Redcay & Courchesne, 2005). The postnatal stage-restricted nature of the deviated brain growth could be due to maldevelopment of the system for programming the size and/or number of brain cells.

However, there have so far been limited systematic molecular and cellular approaches to the relationships between CNVs and frequent co-occurrence of epilepsy or age-dependent size deviation in discrete brain areas in ASD patients. These clinical features of ASD seem to raise the rationale for exploring the role of CNVs in 1q21.1 or 16p11.2 in epileptogenesis and brain formation, because 1) both 1q21.1 and 16p11.2 CNV carriers exhibit microcephaly and macrocephaly (Brunetti-Pierri *et al*, 2008; Mefford *et al*, 2008; Bernier *et al*, 2016; Steinman *et al*, 2016) and 2) the prevalence of seizures is higher in 1q21.1 (deletion (9.5-30%); duplication (6.7-36.4%)) (Brunetti-Pierri *et al*, 2008; Mefford *et al*, 2008) and 16p11.2 (deletion (18%); duplication (20%)) (Steinman *et al*, 2016) CNV carriers than in the general population (≃1%) (World Health Organization, 2019).

Here, to obtain novel insights into the cellular and molecular mechanisms underlying the epilepsy and aberrant brain morphology in ASD, we have investigated the effects of chromosome modification with 1q21.1 deletion (1q del) and duplication (1q dup) on the brain size, differentiation states, transcriptome features, and electrical activities by human neural precursor cell (NPC) organoid model system and 2-dimensional (2-D) neurons. We chose the engineering of the 1q21.1 locus, because no animal or human cell models with 1q dup have yet been generated while mouse models with 16p11.2 deletion and duplication are available (Bult *et al*, 2019). The prevalence of ASD is 5-11% and 13-50% in 1q del and 1q dup, respectively (Brunetti-Pierri *et al*, 2008; Mefford *et al*, 2008; Bernier *et al*, 2016), which are much higher than in the general population (0.6-2.6%) (Maenner *et al*, 2020; Lyall *et al*, 2017).

To this end, we have constructed human ES cell (hESC) clones of both 1q del and 1q dup using a next generation chromosome engineering technique based on CRISPR/Cas9 system (Ran *et al*, 2013), which are theoretically identical with each other except for the target CNV regions. Our target region (840 kb) for 1q del and 1q dup was designed for the distal area of 1q21.1 (Class I deletion/duplication), which harbored all coding genes in the earliest clinical reports of 1q21.1 CNV (Brunetti-Pierri *et al*, 2008; Mefford *et al*, 2008); this area includes the core region among most of 1q21.1 CNV patients with various diseases including ASD (Bernier *et al*, 2016; Pang *et al*, 2020) and corresponds to the region deleted in a mouse model previously established (Nielsen *et al*, 2017).

## Results

### Generation of hESCs with reciprocal 1q21.1 CNVs

In order to induce non-allelic homologous recombination (NAHR) and obtain the reciprocal deletion and duplication clones efficiently, we referred to single-guide CRISPR/Cas9 targeting of repetitive elements (SCORE) method (Tai *et al*, 2016). But since 1q21 region had multiple blocks of highly homologous segmental duplications (SDs), we designed each sgRNA not within but beside two flanking SDs (NBPF11 and NBPF12) by dual guide RNA strategy (Mandal *et al*, 2014) to avoid off-target effects. We transfected Cas9-sgRNA expression vectors into hESCs, called KhES-1, followed by single cell cloning using the limitation dilution method (Figure 1A). First, we performed genomic PCR to detect the CRISPR target site; for deletion, we checked the genomic region flanking the CRISPR target site; for duplication, we amplified the tandem repeat region containing 3’ down sequence followed by 5’ up sequence (Figure 1B). To confirm whether hESCs retained expected CNV, we assessed gene copy numbers of two harboring genes, BCL9 and GJA8, by droplet digital PCR (ddPCR) (Figure 1C). Finally, we performed an array-based comparative genomic hybridization (aCGH) assay to analyze on- and off-target CNVs (Figure 1D and Supplementary Figure 1A and B). We also assessed the mRNA levels of selected genes spanning the 1q21.1 CNV by quantitative real-time-PCR (RT-qPCR) (Supplementary Figure 2). In parallel, we prepared control cells (CTRL) that transfected CRISPR/Cas9 vector without sgRNA.

**Figure 1.**
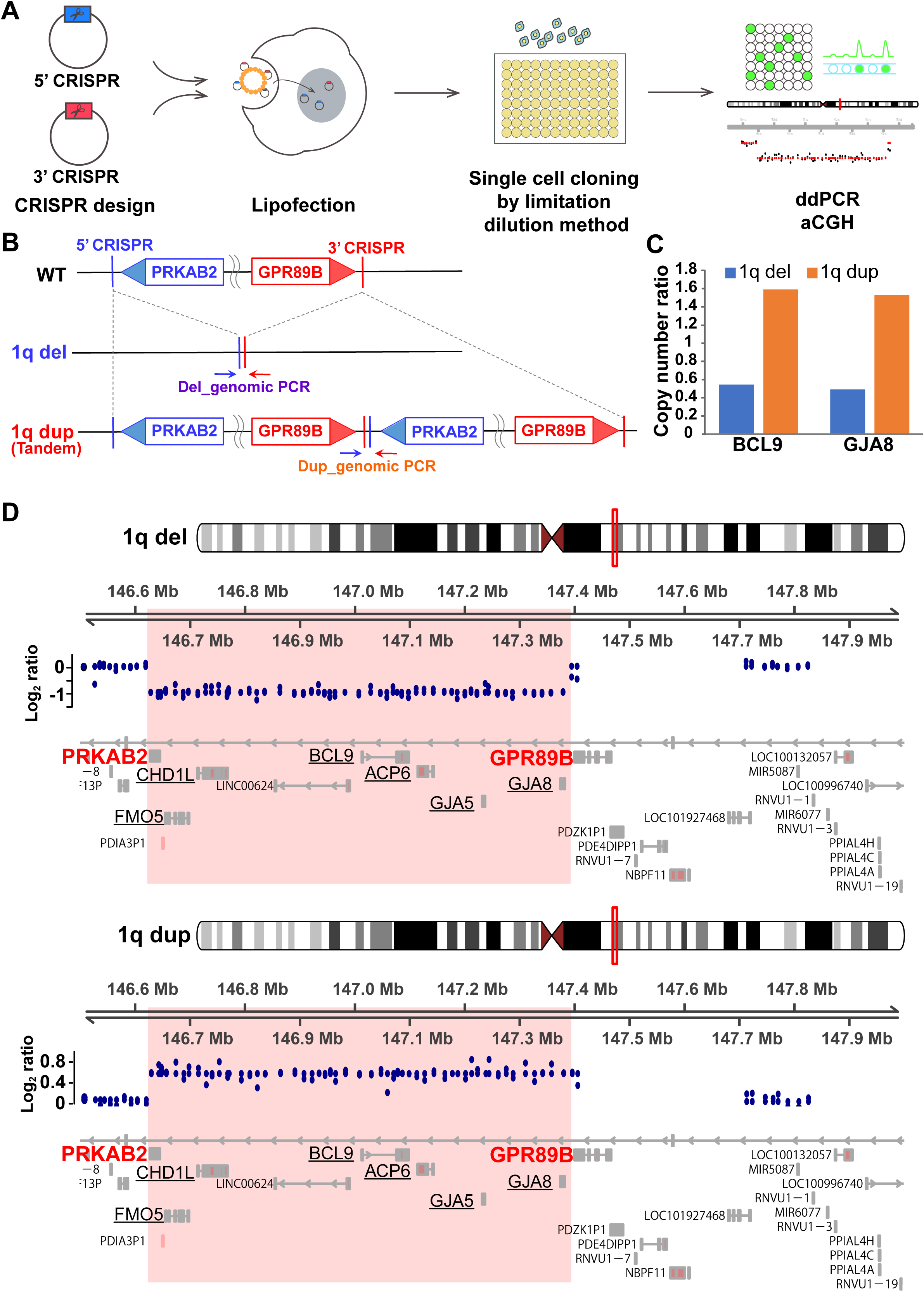
Generation of 1q21.1 mutant human ES cells by CRISPR/Cas9 system. (A) Schematic procedures for construction of 1q21.1 CNV model. (B) Illustration of the targeted segments for 1q21.1 deletion (1q del) and duplication (1q dup) in the 1q21.1 region. Arrows facing each other indicate primer pairs for genomic PCR. (C) ddPCR analysis showing objective CNVs of two targeted genes (BCL9 and GJA8) in each of 1q del and 1q dup candidate clones. Absolute gene copy number of candidate genes were normalized by an endogenous reference gene, AVPR1B (See Materials and Methods). (D) aCGH assays showing objective losses or gains of copy numbers in 1q del and 1q dup candidate clones, respectively. Normalized log_2_ ratio data appear as scatter plots. Pink squares represent the target region. Mb, Megabase.

### Dosage dependent morphometric changes in cortical organoids

We differentiated hESCs into NPC organoids and observed morphological characteristics. We referred to and combined previously described protocols of human cortical organoids (Trujillo *et al*, 2019; Pasca *et al*, 2015; Lancaster & Knoblich, 2014) (Figure 2A). In the early stages, there were no significant differences among the 3 genotypes (Figure 2B and C). On day 17, the circumference size of 1q del NPC organoids was significantly smaller than CTRL but 1q dup was not (p<0.0001 and p=0.097 respectively; Figure 2B and C). However, on day 27, 1q del and 1q dup NPC organoids were significantly smaller (p<0.0001) and or larger (p<0.0001) compared with CTRL, respectively (Figure 2B and C). Next, we analyzed each single-cell in an organoid to answer whether these differences were caused by the number of cells per each organoid or the cell size itself. We made cryosections of 3 genotypes on day 27 and measured the long axis of each cell (Figure 2D), and found no significant differences among them (Figure 2E). Thus, dosage dependent size difference in NPC organoids was caused by the number of cells within an organoid, not by the size of each cell.

**Figure 2.**
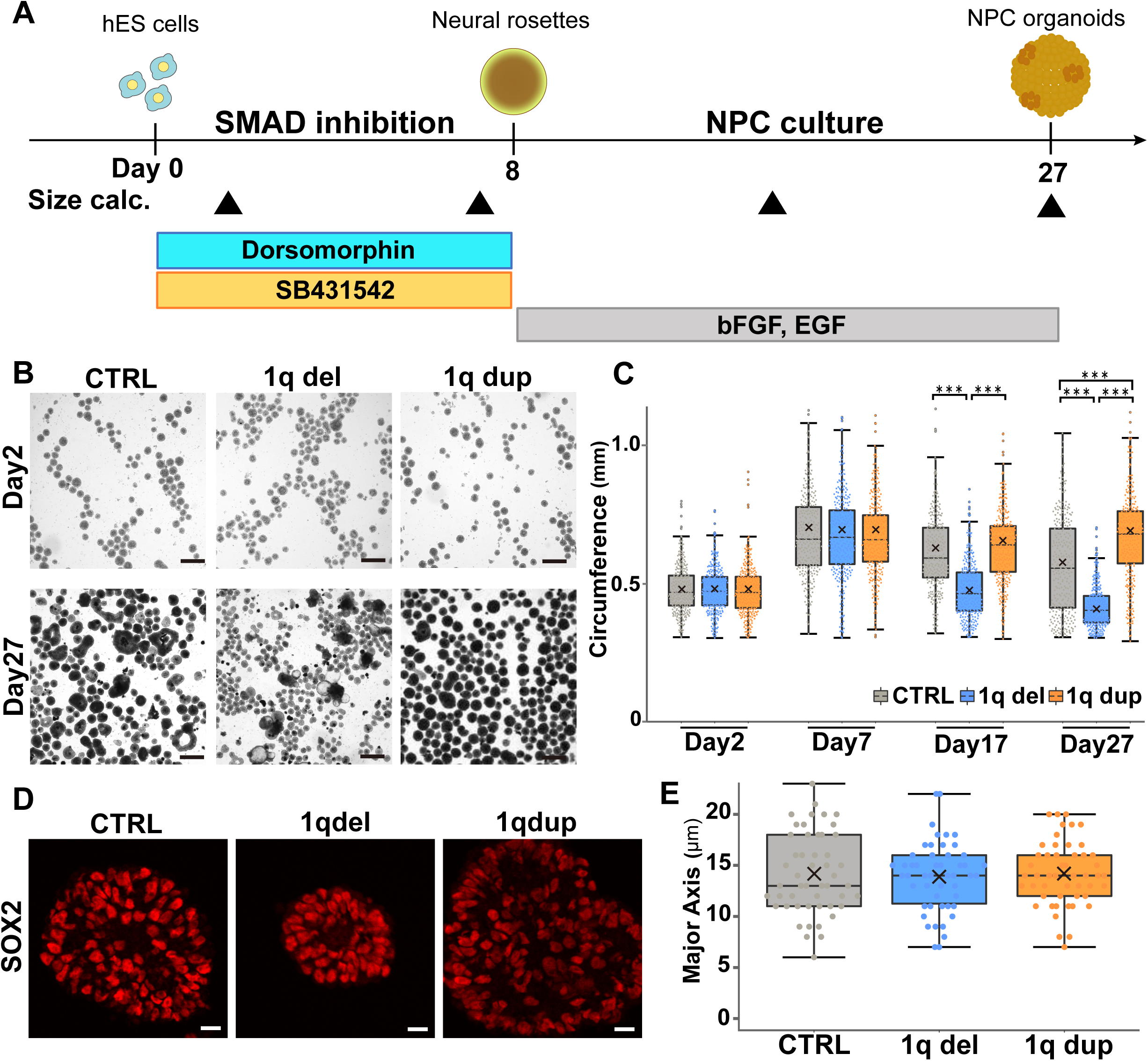
Dosage dependent effects on NPC organoids. (A) Overview of NPC organoid differentiation protocol. Arrowheads indicate the timings of size calculation (day 2, 7, 17, and 27). (B) Representative brightfield images of NPC organoids at day 2 and 27. Each spherical object represents an NPC organoid. Scale bar, 500 µm. (C) Quantitative data of each NPC organoid size showing reciprocal differences of 1q del and 1q dup over time. Sample data of each genotype were randomly extracted from all organoids in the same batch. Data are shown as bee swarm and box plots (n = 300). Each cross represents mean; ***p<0.001; two-tailed Welch’s t-test with Bonferroni correction as post-hoc analysis. Statistical significances were replicated among three independent experiments. (D) SOX2 staining of an NPC organoid to evaluate cellular sizes at day 27. Scale bar, 20 µm. (E) Quantitative data of the major axis length of each cell at day 27 showing no significant differences. Sample data were randomly extracted from 5 organoids per genotype. Data are shown as mean ± SEM (n = 50); two-tailed Welch’s t-test with Bonferroni correction as post-hoc analysis.

### 1q del and 1q dup hESC-derived cortical organoids show reciprocal mature levels

We investigated whether reciprocal 1q21 CNVs affect differentiation pattern and gene expression profile by immunohistochemistry (IHC) and RT-qPCR. The culture protocol of NPC organoids was as described above (Figure 3A). On day 27 of NPC organoid culture, 1q del organoids showed the highest signal intensities of TBR2 (intermediate progenitor marker; also known as EOMES) and TUBB (neuronal marker) among 3 genotypes, whereas 1q dup showed almost no signals in either TBR2 or TUBB (Figure 3B). In addition, RNA expression patterns of NPC organoids on day 25 showed similar results; TBR2 was 1.4 times higher in 1q del and 0.6 times lower in 1q dup than CTRL (1q del, p=0.0086; 1q dup, p=0.0026; Figure 3C). Moreover, 1q del showed 2.3 times higher expression of immature neuronal marker (NCAM1) and 4.0 times higher expression of mature neuronal marker (MAP2) than CTRL (p<0.0001 and p<0.0001 respectively; Figure 3C). We then analyzed gene expression in more mature organoids (Figure 3A). On day 58 (30 days after Matrigel culture), the expressions of both TBR2 and CTIP2 (lower layer cortical marker) were much lower in 1q dup (Figure 3D). Taken together, the neuronal maturity of 1q del seemed to be accelerated, whereas 1q dup decelerated from the NPC stage.

**Figure 3.**
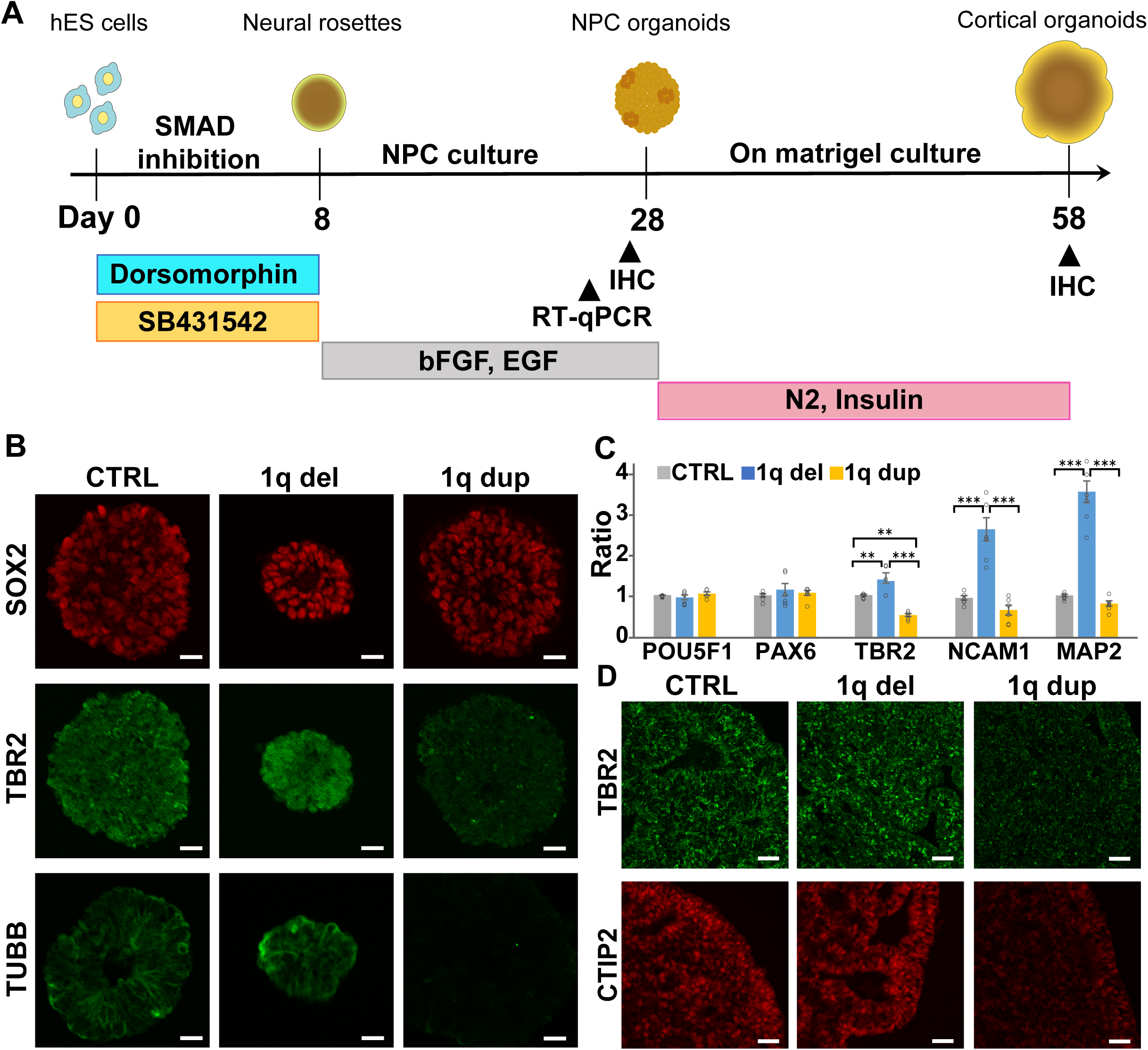
Characteristics of cell differentiations in different genotypes of cortical organoids. (A) Overview of cortical organoid differentiation protocol. Arrowheads indicate the timings of immunohistochemistry (IHC) (day 27 and 58) and RT-qPCR (day 25). (B) Representative images of IHC for proliferative progenitors (SOX2), intermediate progenitors (TBR2) and immature neurons (B-tubulin, TUBB) in organoid cryosections at day 27 showing the highest signals of TBR2 and TUBB in 1q del whereas the lowest in 1q dup. Scale bar, 20 µm. (C) Relative RNA expressions of markers for pluripotent cells (POU5F1), proliferative progenitors (PAX6), intermediate progenitors (TBR2), immature neurons (NCAM1), and mature neurons (MAP2) in organoids at day 25 in accordance with IHC results. Data are shown as mean ± SEM (n=6); **p<0.01, ***p<0.001; one-way ANOVA with Tukey-Kramer test as post-hoc analysis. (D) IHC images for intermediate progenitors (TBR2) and layer V cortical neurons (CTIP2) in organoid cryosections at day 58 showing the lowest signals of both markers in 1q dup. Scale bar, 20 µm.

### Both 1q del and 1q dup hESC-derived neurons have similar features in intrinsic excitability

Although we identified reciprocal phenotypes of 1q del and 1q dup with regard to the organoid size and neural maturity, 1q21.1 CNV carriers also have common clinical features such as epilepsy(Brunetti-Pierri *et al*, 2008; Mefford *et al*, 2008; Bernier *et al*, 2016). Epileptic seizures are generally thought to reflect the hyperexcitability of neuronal networks regardless of underlying factors, e.g., dysfunctions of various ion channels, decreased levels of gamma-aminobutyric acid (GABA) and glutamic acid decarboxylase (GAD) in epileptic focus, and hyperactivation of glutamatergic receptors (Chang & Lowenstein, 2003; Badawy *et al*, 2009). Thus, we next examined the spontaneous electrical activity of hESC-derived 2-D neurons by using multi-electrode array (MEA). Referring to a published protocol (Fujimori *et al*, 2017), we cultured neurospheres of 1q del, 1q dup and CTRL, then we dissociated these cells onto microelectrode arrays (Figure 4A and B). Spontaneous spike activities began to increase from around 21 days post-dissociation (21 dpd) (Figure 4C and D and Supplementary Figure 3A). At 28 dpd, both 1q del and 1q dup 2-D neurons showed significantly higher frequency of spontaneous spikes than the CTRL (p<0.0001; Figure 4E). Moreover, we observed many burst firings in some of 1q del and 1q dup neurons at 35 dpd, and 1q del neurons showed significantly more bursts compared with CTRL (1q del, p=0.0019; 1q dup, p=0.16; Supplementary Figure 3B). These results suggest that 1q del and 1q dup neurons show similar electrical properties with a little higher degree of intrinsic excitability in 1q del.

**Figure 4.**
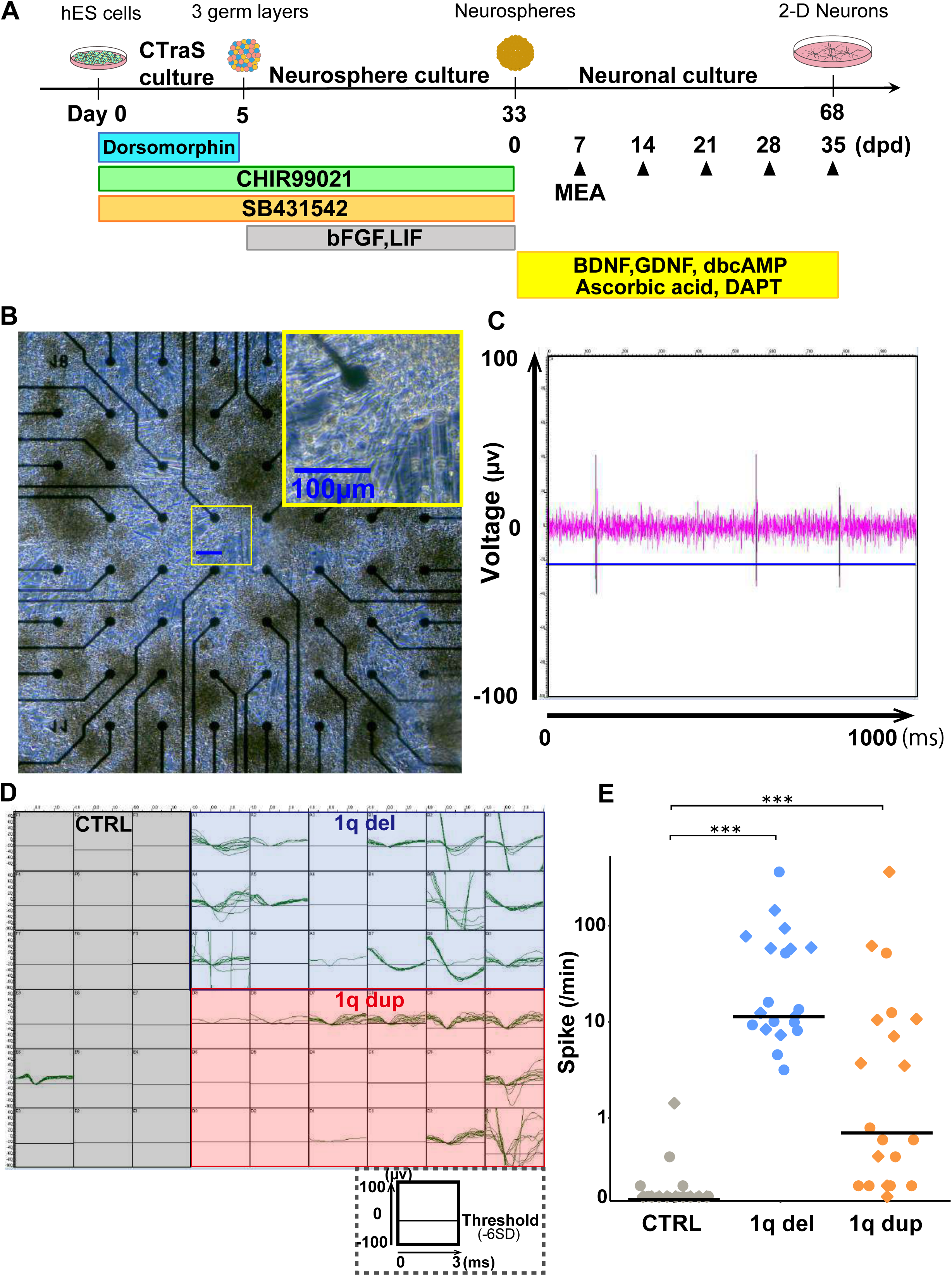
Multi-electrode array (MEA) analysis by 2-dimensional differentiated (2-D) neurons. (A) Overview of 2-D neuron culture protocol. After adherent culture with 3 chemical inhibitors (Dorsomorphin, CHIR99021 and SB431542), cells were kept in suspension from day 5 to day 33 and then dissociated onto MEA array dishes. Arrowheads represent the timings of MEA analysis (7, 14, 21, 28, and 35 days post-dissociation (dpd)). CTraS, chemically transitional embryoid-body-like state (Fujimori et al., 2017). (B) Representative brightfield images of cultured neurons at 28 dpd. Each black dot represents an electrode. Yellow square at the top-right corner is the magnified image showing many neurites. Scale bar, 100 µm. (C) Representative waveform of spontaneous spikes. Sharp signals which go across the threshold (blue horizontal line) are detected as spikes. (D) Overlay images of spontaneous electrical activities for 5 minutes at 28 dpd. Each square represents one electrode. Both 1q del and 1q dup had many more spikes than CTRL. (E) Median spike frequency per electrode showing much higher frequency in both mutants compared with CTRL. Top 10 data from 2 batches were extracted; circle dots and square dots represent each batch. Data are shown as bee swarm plots (n = 20); ***p<0.001; Mann-Whitney U test with Bonferroni correction as post-hoc analysis.

### Cellular profiles of reciprocal CNVs show clear differences in the speed and direction of cell differentiation

Although 1q21.1 CNV showed reciprocal changes in the organoid size and mature level, and showed common features of intrinsic neural excitability, it was still unclear how differentiation patterns were affected by the CNV. To investigate cellular diversity and expression profiles during early development, on day 27 we performed single-cell RNA sequence analysis (scRNA-seq) using NPC organoids. We first performed k-means clustering on the dataset of combined 32,171 cells (1q del; 10,682, 1q dup; 11,987, CTRL; 9,502) to identify clustering patterns and the breakdown by genotypes. Based on the expression patterns of marker genes, we finally set 8 clusters: neural stem cell (’NSC’, the most immature); neural progenitor cell-1 (’Progenitor-1’); ’Progenitor-2’ (more mature than Progenitor-1); glial cell including precursors of astrocytes and oligodendrocytes (’Glial’); endothelial cell (’Endothelial’); intermediate progenitor cell (’IPC’); GABAergic neuron (’GABAergic’); and unspecified cell (’Other’) (Figure 5A-C). Distribution patterns of 1q del, 1q dup and CTRL cells were shown in Uniform Manifold Approximation and Projection (UMAP) (Figure 5A). Approximately 60% of 1q dup cells (7,300 cells) belonged to the Glial cluster, while only 81 cells of 1q del were included in this cluster. In contrast, approximately 70% of 1q del cells belonged to Progenitor-1 or Pprogenitor-2 cluster (5,208 cells and 2,409 cells, respectively), both of which contained small numbers of 1q dup cells (17 cells and 1 cell, respectively). Despite being quite a small cluster, more than 90% of the cells in the GABAergic cluster were derived from 1q del (188 cells). Thus, scRNA-seq data showed 1q del and 1q dup NPC organoids contained different cellular compositions. The fact that GABAergic cells were almost exclusively derived from 1q del (although still very limited) may indicate the difference in a mature level, which was consistent with gene expression analysis data shown below.

**Figure 5.**
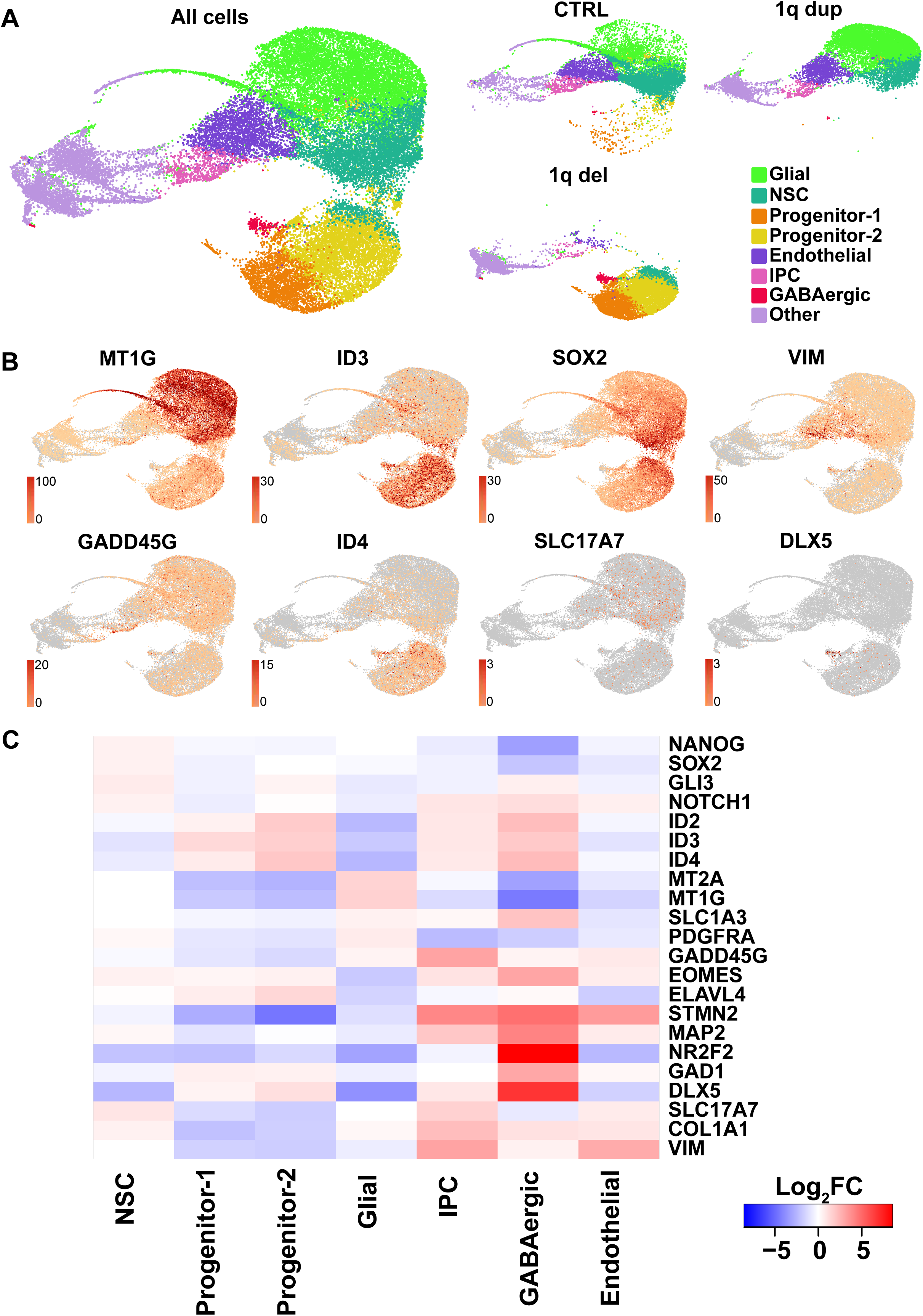
Gene expression profiles of human differentiated NPC organoids by single cell RNA sequencing (scRNA-seq). (A) UMAP plots of the integrated datasets. The left-side plot shows whole samples and right-side plots show separate data by different genotypes. Yellowish green as glial cell (Glial); blue green as neural stem cell (NSC); orange as neural progenitor cell-1 (Progenitor-1); yellow as Progenitor-2; dark purple as endothelial cell (Endothelial); pink as intermediate progenitor cell (IPC); red as GABAergic neuron (GABAergic); and light purple as unspecified cell (Other). (B) Feature plots show expression levels of cell-type-specific markers: MT1G; glial, ID3; progenitor-1, SOX2; NSC, VIM; endothelial, GADD45G; IPC, ID4; progenitor-2 (relatively mature than ID3), SLC17A7; excitatory neuron, DLX5; inhibitory neuron. Coloring of the plots represents normalized gene unique molecular identifier (UMI) counts. (C) Heatmap shows cluster-specific gene expression across cell clusters. Coloring represents log_2_ ratio of the normalized gene expressions in each cluster relative to all other clusters for comparison.

### Enrichment analyses of gene expression datasets in 1q del and 1q dup NPC organoids

We next performed enrichment analysis by using scRNA-seq dataset. We extracted differentially expressed genes (DEGs) between each CNV versus CTRL if the following conditions were satisfied; p < 0.05 and |log_2_FC| >1.2. In bulk analysis, the number of upregulated DEGs compared with CTRL (up-DEGs) was 560 and 85 in 1q del and 1q dup, respectively, and the number of common up-DEGs between 1q del and 1q dup was 17. On the other hand, the number of downregulated DEGs (down-DEGs) was 701 and 281 in 1q del and 1q dup, respectively, among which 32 DEGs were common (Figure 6A).

**Figure 6.**
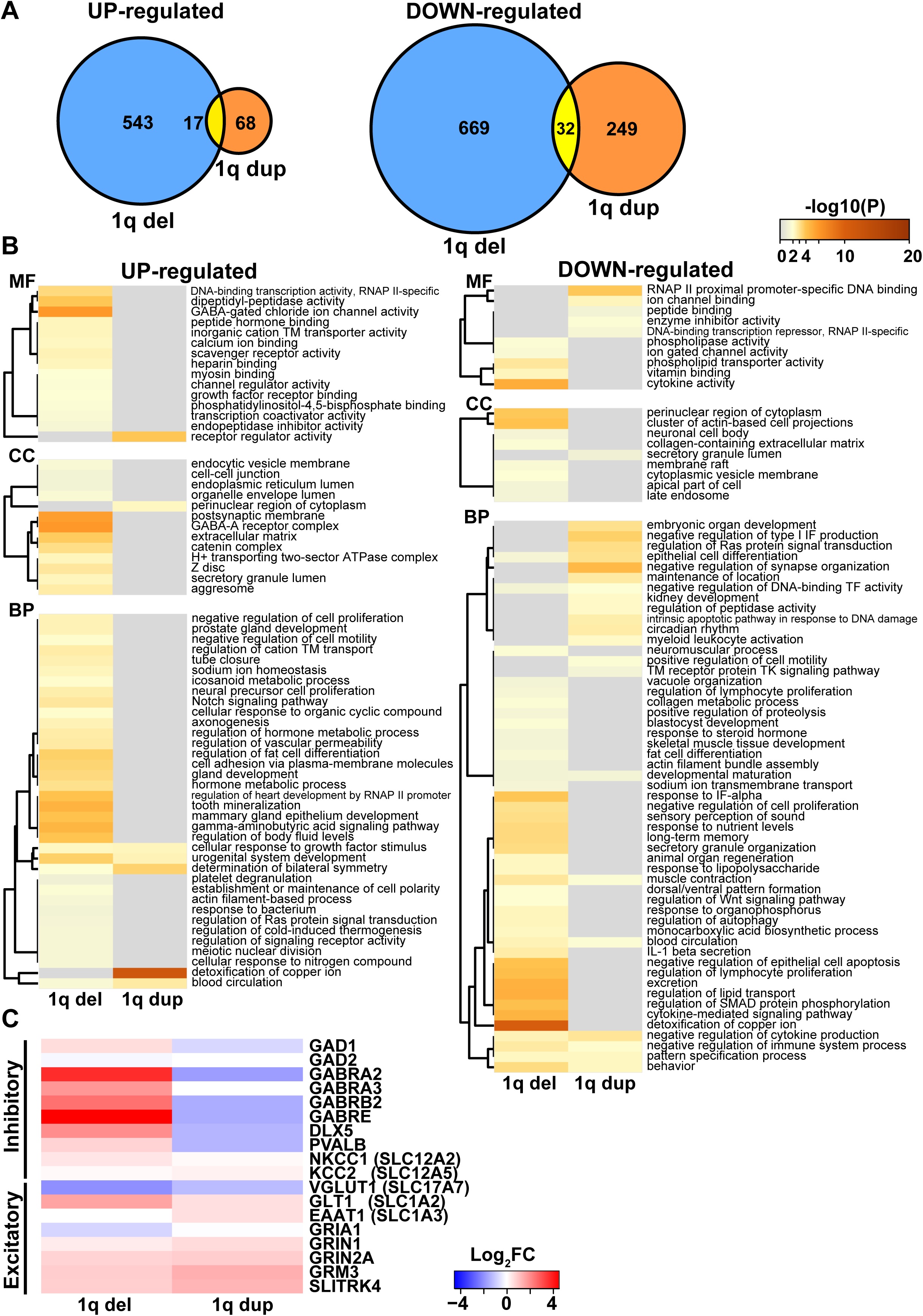
Enrichment analysis of NPC organoids. (A) Venn diagrams of differentially expressed genes (DEGs) by enrichment analysis in bulk. Left-side graph shows up-regulated (up-) DEGs and right-side shows down-regulated (down-) DEGs compared with CTRL. Few DEGs were overlapped between 1q del and 1q dup in both up- and down-DEGs. (B) Heatmap of gene ontology (GO) terms enrichment analysis in the NSC cluster showing quite different annotations between 1q del and 1q dup. Left-side and right-side maps show GO terms by up-DEGs and down-DEGs, respectively, compared with CTRL. Coloring represents log_10_ p-score of each GO term. MF, molecular function; CC, cellular component; BP, biological process. (C) Heatmap of gene expressions of inhibitory and excitatory neuron markers in the NSC cluster. Coloring represents log_2_ ratio of the normalized gene expressions in each mutant relative to CTRL. Expressions of inhibitory markers were higher in 1q del whereas lower in 1q dup compared with CTRL. On the other hand, expressions of excitatory markers showed similar altered patterns in both mutants.

#### (1) Analyses with Disease Ontologies

It is well documented that 1q del and 1q dup carriers exhibit one or more symptoms observed in not only neuropsychiatric disorders but also various diseases in peripheral organs such as eye, kidney, heart and muscle (Brunetti-Pierri *et al*, 2008; Pang *et al*, 2020; Bernier *et al*, 2016). Hence, we performed enrichment analyses on diseases. In agreement with those reports, neuropsychiatric terms such as ASD, epilepsy, schizophrenia, mood disorders and abnormal behavior, as well as disease terms in peripheral organs, e.g. eye, heart, kidney and muscle, were enriched in both 1q del and 1q dup (Supplementary Table 1) although most DEGs were not overlapped in 1q del and 1q dup as described above (Figure 6A). Even when we focused on common DEGs, most gene functions correlated with either brain or these peripheral organs (Supplementary Table 2). Nevertheless, there were some characteristic differences; DEGs in 1q del contain terms related to schizophrenia symptoms e.g., delusion, anhedonia and impaired cognition (Supplementary Table 1). This result seems concordant with the fact that schizophrenia is more relevant in 1q del than 1q dup; frequency and odd ratio (OR) of each CNV in schizophrenia patients have been reported as 0.17-0.23% (OR 8.1-14.8) and 0.13-0.14% (OR 3.5-4.2), respectively(Malhotra & Sebat, 2012; Stefansson *et al*, 2008; Marshall *et al*, 2017).

#### (2) Gene Ontology (GO) analyses by NSC cluster

For further analysis, we mainly focused on the NSC cluster for the following reasons: firstly, cells from all 3 genotypes were affiliated in this cluster enough to extract sufficient numbers of DEGs; secondly, this cluster was thought to be appropriate for identifying heterogeneity in the very early stage of cell differentiation. We performed GO analysis using Metascape (Zhou *et al*, 2019) and Enrichr (Kuleshov *et al*, 2016). We input DEGs of 1q del and 1q dup described above and analyzed molecular function (MF), cellular component (CC), and biological process (BP) (Figure 6B). Among up-DEGs in 1q del, we found GABA-related terms in all 3 categories, such as ’GABA-gated chloride ion channel activity’ (MF), ’GABA-A receptor complex’ (CC), and ’gamma-aminobutyric acid signaling pathway’ (BP). We also found neuronal GO terms such as ’post synaptic membrane’ (CC) and ’axonogenesis’ (BP) in 1q del, whereas GO terms associated with epithelial differentiation, developmental maturation and regulation of synapse organization were found in down-DEGs of 1q dup. Regarding pathway-related terms in 1q del, JAK2/STAT3, which is associated with neural protection and proliferation, and GABA-related pathways such as nicotine and morphine addiction were upregulated, whereas terms related to cell cycle progression were downregulated (Supplementary Table 3). On the other hand, there were many up-regulated pathways associated with cell proliferation, i.e., MAPK, PI3K-AKT, TGF-beta and RAP1 pathways in 1q dup. There were almost no GO terms related to mature neuronal cells in 1q dup, which may suggest that they were resistant to neural differentiations. These results suggest that 1q del had already started to differentiate into neurons in this period, meanwhile, 1q dup still maintained its stem cell state.

#### (3) Analyses with GABAergic and glutamatergic neurotransmission-related gene sets

Since we found enrichment in GABA-related terms of 1q del described above, we further checked expressions of GABAergic and glutamatergic related genes. Most GABA-related genes were enriched in 1q del, whereas they were mostly downregulated in 1q dup (Figure 6C). As for cation-Cl^-^ co-transporters (CCCs), which are related to Cl^-^ homeostasis, Na^+^ -K^+^-2Cl^-^ co-transporter 1 (NKCC1) was upregulated in 1q del, while K^+^-2Cl^-^ co-transporter 2 (KCC2) had almost no change compared with CTRL. These expression patterns of CCCs indicated that GABA acted in an excitatory manner (Watanabe & Fukuda, 2015). With respect to glutamatergic genes, both 1q del and 1q dup showed similar altered expressions compared with CTRL (Figure 6C). These results indicate that 1q del cells are in a more mature state with more GABAergic components, whereas 1q dup cells are in a more undifferentiated and proliferative states with more glutamatergic components.

## Discussion

In this study, we constructed reciprocal hESC models of CNV in distal 1q21.1 region and analyzed cellular properties using differentiated cells. This is the first study analyzing reciprocal 1q21.1 CNV human cell models and the first human ES model of CNV. Although many investigations have been conducted on neurodevelopmental disorders using induced pluripotent stem cells (iPSCs) (Lee *et al*, 2020), in this study we used ESCs because they are deemed to have advantages compared with iPSCs. Genetic and epigenetic alterations including increased CNVs have been uncovered in iPSCs (Panopoulos *et al*, 2011; Hussein *et al*, 2011; Gore *et al*, 2011; Lister *et al*, 2011; Laurent *et al*, 2011) and ESCs are still the gold standard for in vitro pluripotency (Bilic & Izpisua Belmonte, 2012). In addition, our isogenic cell models are theoretically identical to each other except for the target region, therefore these are ideal models for investigating how CNVs affect cellular phenotypes.

To induce NAHR as much as possible in our target region of the distal area of 1q21.1, we designed each sgRNA beside two flanking SDs (NBPF11 and 12). The proximal 1q21.1 region is known to be associated with an independent syndrome named Thrombocytopenia Absent Radius (TAR) syndrome (Pang *et al*, 2020; Klopocki *et al*, 2007). We only focused on distal 1q21.1 region although some clinical patients have longer mutations overlapping both proximal and distal regions (Brunetti-Pierri *et al*, 2008; Mefford *et al*, 2008). Thus, our target region covered the core regions of distal 1q21.1 CNV (Pang *et al*, 2020).

Using these reciprocal hESC models of distal 1q21.1 CNV, we cultured cortical organoids and identified dosage-dependent morphological changes that may underlie the reciprocal brain size observed in CNV carriers. Size measurement of each cell in an organoid revealed that it originated from the cell number, not from the size of each cell itself, which suggested the difference of proliferation status between 1q del and 1q dup. We also identified reciprocal mature levels in cortical organoids from IHC, RT-qPCR and scRNA-seq. These results indicate that 1q del organoids tend to differentiate into neurons, whereas 1q dup maintain the undifferentiated state.

In addition to reciprocal changes, 1q del and 1q dup had some common features. MEA analysis to evaluate neural electrical activity using hES-derived 2-D neurons revealed that both CNVs exhibited hyperexcitability compared with CTRL. In addition, GO enrichment terms from common DEGs identified comorbidity phenotypes of both 1q21.1 CNV carriers, such as cataract and cardiovascular disease (Brunetti-Pierri *et al*, 2008; Bernier *et al*, 2016; Pang *et al*, 2020). Thus, our data using isogenic cell lines recapitulate some elements of the clinical features found in 1q21.1 CNV carriers. These results suggest that clinical symptoms in 1q21.1 CNV carriers are derived from the causative mutations rather than other factors such as latent genetic background or environmental factors.

### Dosage-dependent morphological changes

We observed reciprocal changes in the size of NPC organoids. We assumed several factors are involved in this phenomenon. Firstly, we focused on genetic factors harboring our target region. The target region contains 8 coding genes: PRKAB2, FMO5, CHD1L, BCL9, ACP6, GJA5, GJA8, and GPR89B. Among these genes, PRKAB2 plays a role in homeostasis of glucose and lipid metabolism, and its knock out (KO) condition reduced ghrelin-induced neuroprotective effect (Bayliss *et al*, 2016). This may be related to the decreased cell number in organoids of 1q del. CHD1L and BCL9 are members of oncogenes. Since several genetic studies have shown extensive overlap in risk genes for autism and cancer (Crawley *et al*, 2016), these genes would be possible candidates for 1q21.1 pathogenesis. A previous paper reported that CHD1L, a chromatin remodeling factor, induced differentiation of hESCs into PAX6-positive NPCs (Dou *et al*, 2017), it also plays a role in tumor proliferation and progression (Liu *et al*, 2016; Chen *et al*, 2009b). Thus, CHD1L may cause dosage-dependent proliferation of 1q21.1 dup NPCs. BCL9 is associated with Wnt-driven tumorigenesis by stabilizing beta-catenin (Gay *et al*, 2019). Since Wnt/beta-catenin signaling induces NPC proliferation in the early developmental stage (Chenn & Walsh, 2002; Hirabayashi *et al*, 2004), it might be a factor for dosage-dependent effects as well.

Secondly, we focused on the results of the signaling pathway. Our data showed the up-regulation of MAPK and PI3K-AKT pathways in 1q dup NPC organoids. Both RAS/MAPK and PI3K/AKT/mTOR are well known pathways important for cell growth, differentiation and synaptic plasticity (Borrie *et al*, 2017). Gene mutations in the RAS/MAPK pathway cause developmental disorders called RASopathies; they have overlapping symptoms, e.g. macrocephaly, cognitive and behavioral impairments, craniofacial dysmorphism, cutaneous lesions, cardiovascular problems, and increased tumor risk. On one hand, genetic mutations in PI3K/AKT/mTOR pathways are associated with hamartoma syndromes that are characterized by brain overgrowth, neurological problems and benign tumors (Borrie *et al*, 2017; Dobyns & Mirzaa, 2019). These results suggest that altered gene expressions in these two pathways may underlie excessive cell proliferation in 1q dup organoids. To verify dosage-dependent effects in cell differentiation patterns, it is desirable to get additional data at different time points in the future.

### Hyperexcitability and seizures

Seizures are common features in both 1q del and 1q dup, and the prevalence in both CNVs is much higher than in general population (Brunetti-Pierri *et al*, 2008; Mefford *et al*, 2008; World Health Organization, 2019). Our MEA analysis data showed higher spontaneous spikes in both 1q del and 1q dup 2-D neurons compared with CTRL. In addition, 1q del neurons showed significantly higher burst frequency than CTRL. Furthermore, scRNA-seq data showed that both 1q del and 1q dup NSCs had altered expressions of glutamatergic components compared with CTRL. Generally, epileptic seizures are thought to be generated by the imbalance between excitatory and inhibitory (E/I) signals (Chang & Lowenstein, 2003; Badawy *et al*, 2009). The majority of epileptic patients show interictal spikes (Smith, 2005; Staley & Dudek, 2006); repetitive action potentials are driven by sustained glutamate release in epileptic focus, which may contribute to epileptogenesis (Staley & Dudek, 2006). Thus, hyperexcitability and altered glutamate systems in our data may suggest the neuronal E/I imbalance underlying the high prevalence of epilepsy in 1q21.1 CNV carriers. However, there are some limitations in our experiments. Firstly, the 2-D neurons used in this study were rather immature because their neuronal subtypes were not well determined. Secondly, we only studied spontaneous neuronal activity in MEA analysis; hyperreactivity to external stimuli were not analyzed this time. To verify the dysfunction of glutamatergic system, further experiments such as pharmacological tests would be needed.

### Alterations of GABAergic components in 1q del cells

Our scRNA-seq data showed that 1q del cells had the highest expressions of GABAergic components among 3 genotypes. These results indicate two possibilities; one possibility is that it simply reflected a different developmental stage. A previous paper showed that expression of GABAergic components was mainly restricted to cortical organoids at the later stage (Trujillo *et al*, 2019), which were concordant with in vivo study (Uylings *et al*, 2002). Another possibility is the GABAergic dysfunction in 1q del. It is known that alterations of GABAergic components are related to some diseases such as epilepsy (Chang & Lowenstein, 2003; Badawy *et al*, 2009) and schizophrenia (de Jonge *et al*, 2017; Lewis *et al*, 2005), both of which are observed in 1q del carriers(Malhotra & Sebat, 2012; Brunetti-Pierri *et al*, 2008; Mefford *et al*, 2008; Stefansson *et al*, 2008; Marshall *et al*, 2017).

It seemed contradictory that 1q del 2-D neurons exhibited hyperexcitability while 1q del NPCs showed high expression of GABAergic components in our results. We suggest a possible reason stemming from their developmental stages. 1q del NPCs showed up-regulation of NKCC1 while KCC2 was almost unchanged compared with CTRL. NKCC1 plays an important role in increasing intracellular Cl- concentration such that GABA acts as an excitatory transmitter in NPCs and immature neurons (Watanabe & Fukuda, 2015; Payne *et al*, 2003; Koyama *et al*, 2012). On the other hand, KCC2 works as a Cl^-^ extruder and up-regulation of this molecule results in hyperpolarizing action of GABA in mature neurons (Watanabe & Fukuda, 2015; Payne *et al*, 2003; Ganguly *et al*, 2001). Thus, roles of GABA switches dramatically along with development.

Down-regulation of GAD67 and up-regulation of GAT in presynaptic parvalbumin-positive (PV) neurons, and up-regulation of GABAergic receptors (especially alpha 2 subunit) in axon initial segments of postsynaptic pyramidal neurons are the most common findings in postmortem studies of schizophrenia patients (de Jonge *et al*, 2017; Lewis *et al*, 2005); this expression pattern is similar to our results. Although it is still unknown whether GABRA2 upregulation is the initial excessive GABAergic activity or compensatory reaction for the presynaptic reduction of GABA, altered expression patterns in 1q del would indicate GABAergic dysfunction. However, since the NPC organoids in our experiments were still immature, a longer follow up is needed to analyze the transition of neural maturation in all genotypes in order to examine the E/I imbalance hypothesis in the forebrain of ASD.

## Conclusion

Our data from the different approaches recapitulate various characteristics in reciprocal 1q21.1 CNV carriers and suggest some mechanisms underlying clinical features. Our findings shed light on the importance of NPC as a regulator of proliferation and a determinant of neural fate in 1q21.1 CNV. We also found that neuronal hyperexcitability is a common feature of reciprocal 1q21.1 CNVs despite their different expression patterns. Our cell models of 1q21.1 CNV could be applied as tools to further explore the genomic basis of comorbidities and the onset of ASD. Extending our knowledge into other genetic models would deepen our understanding of the mechanisms of neurodevelopmental disorders.

## Materials and Methods

### Human embryonic stem cell (hESC) source

A human ES Cell line, KhES-1, were supplied from Institute for Frontier Medical Sciences (Kyoto University, Kyoto, Japan). These cells were cultured using Cellartis DEF-CS 100 Culture System (DEF-CS medium; Takara) under feeder-free condition. Medium was exchanged daily and cells were manually passed every 4-5 days when they became confluent. Pluripotency state was checked by Alkali-phosphatase staining before and after genome editing experiments (Supplementary Figure 4). All experiments were approved by an institutional ethics committee and performed following the hES cell guidelines of the Japanese government.

### Generation of 1q21.1 deletion and duplication CNVs using CRISPR/Cas9 system

To generate target CNVs, we designed two Cas9-sgRNA expression vectors: one for up-stream of 5’ end (PRKAB2) and another for down-stream of 3’ end (GPR89B). We used pX330-U6-Chimeric_BB-CBh-hSpCas9 (Addgene plasmid #42230) and guide RNAs (gRNAs) were designed using CRISPR direct (http://crispr.dbcls.jp/) and CRISPR design (http://crispr.mit.edu:8079/) software to avoid off-target effects. Each gRNA was cloned into the vector using BbsI restriction site. The sequence of each gRNA was as below.

5’ up_gRNA: 5’-AAATCTCTGGTATGGGGTCC-3’, chromosome 1: 147,154,711-147,154,730

3’ down_gRNA: 5’-TCTAGGTATGACTAGAAGTC-3’, chromosome 1: 147,994,573-147,994,592

Guide sequences were confirmed by Sanger Sequencing. Before transfection, all plasmids were purified using QIAprep Spin Miniprep Kit (QIAGEN) according to the manufacturer’s protocol. We co-transfected gRNA_cloning vectors (1 µg each) and pSpCas9(BB)-2A-Puro plasmid (1 µg) using lipofectamine 3000 reagents (Thermo Fisher Scientific) according to the manufacturer’s instructions. At 2 days after transfection, puromycin selection was performed for 48 hours. At 5 days after transfection, we performed single-cell cloning by limitation dilution method.

### Screening of individual hESC colonies

Confirmation of target CNVs were performed by genomic PCR, ddPCR and aCGH assay.

#### Genomic PCR assay

Genomic DNA (gDNA) was extracted from each well of single cell-derived clones. For detecting deletion, the genomic region flanking the CRISPR target site was amplified; for detecting duplication, the tandem repeat region containing 3’ down sequence followed by 5’ up sequence was amplified (Figure 1B). Primers were synthesized by Integrated DNA Technologies (IDT; Sigma-Aldrich). Sequences of designed primers were as below:

1q del_PCR_F: 5’-AGTTTGGGCCCCGATGAAAT-3’
1q del_ PCR_R: 5’-ACACACACAATGCCCACTGA-3’
1q dup_ PCR_F: 5’-TTGGTGCTCAAGGAACCTGT-3’
1q dup_ PCR_R: 5’-TGCAGCTCTGTGTCTGTCAG-3’

Genotyping PCR were performed using 100 ng of gDNA and LA Taq DNA polymerase (Takara), with the following cycling conditions: 94 °C for 1 min; 98 °C for 10 s, 68 °C for 1.5 min (40 cycles).

#### ddPCR assay

Droplets containing gDNA samples and QX200™ ddPCR™ EvaGreen Supermix were mixed for Automated Droplet Generator (Bio-Rad) according to the manufacturer’s protocol. Then, samples were analyzed using QX200™ Droplet Reader (Bio-Rad). Primers in exon 6 of BCL9 and exon 1 of GJA8 were used for screening. Primers outside of the target region were also designed as an endogenous reference gene in exon 1 of AVPR1B on 1q32.1. Primers were synthesized by IDT with the following sequences:

ddPCR_BCL9_F: 5’ ACACACCACACTCGATGACC 3’
ddPCR_BCL9_R: 5’ GGGCTCTGTGGCAGTAGTTT 3’
ddPCR_GJA8_F: 5’ ACCCTGCTGAGGACCTACAT 3’
ddPCR_GJA8_R: 5’ TACAGAGGCAGGATCCGGAA 3’
ddPCR_AVPR1B_F: 5’ GGCCAAGATCCGAACAGTGA 3’
ddPCR_AVPR1B_R: 5’ GGACACTGAAGAAGGGAGCC 3’

PCR reactions for droplets were performed according to the manufacturer’s instructions with the following cycling conditions: 95 °C for 5 min; 95 °C for 30 s, 60°C for 1 min (40 cycles); 4 °C for 5 min, 90 °C for 5 min. Copy numbers of each targeted gene were normalized by the reference gene. Clones with nearly half scores of wild type (WT) were regarded as 1q del clones, and nearly 1.5 times were regarded as 1q dup clones. Positive clones were expanded and used to extract gDNA samples for further analysis.

#### aCGH assay

aCGH assay for both 1q del and 1q dup clones was performed on the Agilent 4 × 180K SurePrint G3 Human CGH Microarray (design number 022060) according to the manufacturer’s protocol. DNA from WT hESCs was used as reference genome. Analyses were performed as previously reported (Kishimoto *et al*, 2015). We referred to GRCh37/hg19 human reference genome (UCSC Genome Browser, https://genome.ucsc.edu/cgi-bin/hgGateway/).

### Real-time quantitative PCR (RT-qPCR) for hESC clones

Total RNA was extracted using TRI Reagent (Molecular Research Center) according to the manufacturer’s instruction. Reverse transcription reactions were performed with SuperScript IV Reverse Transcriptase (Thermo Fisher Scientific) in 20 µl total volume containing 500 ng of RNA. Primers within the target region were designed in exon 3 of PRKAB2, exon 6 of BCL9, and exon14 of GPR89B. Primers outside the target CNV region (both 5’ upstream and 3’ downstream) and GAPDH as a housekeeping gene were also designed. Primers spanning introns were designed using Universal ProbeLibrary Assay Design Center (https://qpcr.probefinder.com/organism.jsp/). Primers were synthesized by IDT with the following sequences:

qPCR_PRKAB2_ex3_F: 5’ GCAGCAGGATTTGGAGGACT 3’
qPCR_PRKAB2_ex3_R: 5’ CCTTGCCTCCTTCAGACCAG 3’
qPCR_BCL9_ex6_F: 5’ ACACACCACACTCGATGACC 3’
qPCR_BCL9_ex6_R: 5’ GGGCTCTGTGGCAGTAGTTT 3’
qPCR_GPR89B_ex14_F: 5’ TTGTCTCCTCTGTGCTGCTG 3’
qPCR_GPR89B_ex14_R: 5’ GCCATTTGCTTCTCTGGTGC 3’
qPCR_HJV_ex4_F: 5’ CCATGGCCTTCTCAGCTGAA 3’
qPCR_HJV_ex4_R: 5’ TGGTTATAGCTCCCCGACGA 3’
qPCR_NUDT4B_ex1_1_F: 5’ GACCAGTGGATTGTCCCAGG 3’
qPCR_NUDT4B_ex1_1_R: 5’ TCCTTTGACTCCAGCCTCCT 3’
qPCR_GAPDH_F: 5’ ACGGGAAGCTCACTGGCATGGCCTT 3’
qPCR_GAPDH_R: 5’ CATGAGGTCCACCACCCTGTTGCTG 3’

qPCR reactions were carried out in 25 µl total volume containing 5 ng of cDNA sample and 0.3 µM designed primers and SYBR Green Master Mix (Bio-Rad). Amplification was performed by Step One Plus (Applied Bio System) under the following cycling conditions: 50 °C for 2 min, 95 °C for 10 min; 95 °C for 15 s, 60 °C for 1 min (40 cycles). Values were normalized to GAPDH expression. Statistical analysis for copy number was performed using Welch’s t-test followed by Bonferroni correction for multiple comparisons.

### Primary astrocyte culture for Multi-Electrode Array (MEA) analysis

Primary astrocytes were extracted as previously described with some modification (Hosoi *et al*, 2004). Postnatal day 0 (P0) mouse cerebral cortices were dissociated using 0.25 % Trypsin (Gibco). After pipetting about 40 times, these cells were filtered and seeded on 10 cm dishes at a density of cells from 1 mouse per 1 dish then cultured in humidified atmospheres (5 % CO2). Astrocytes were then purified by shaking the culture dish to remove microglia and oligodendrocytes. The medium was exchanged after 3 days and cells were manually passed from 1 dish into 2 dishes after 1 week. Passages were performed several times until cells ceased to proliferate. To prepare for MEA recording, astrocytes were seeded on microelectrode arrays (60MEA200/30iR-Ti and 60-6wellMEA200/30iR-Ti, Multi Channel Systems) coated with 0.05 mg/ml poly-DL-ornithine (Sigma-Aldrich) and 10 µg/ml fibronectin (Sigma-Aldrich) at a density of 2 x 10^5^ cells per array.

### 2-dimensional (2-D) neuronal differentiation culture for MEA analysis

For neuronal differentiation, we followed the protocol as previously published (Fujimori *et al*, 2017) with slight modifications. hESCs were cultured on 1 well of a 6-well plate in induction medium containing DEF-CS medium, 3 µM SB431542 (WAKO), 3 µM Dorsomorphin (WAKO) and 3 µM CHIR99021 (WAKO) for 5 days. The medium was changed daily. On day 5, colonies were detached and dissociated into single cells using TrypLE Select (Life Technologies). These cells were cultured in suspension at a density of 2 x 10^6^ cells per Petri Dish for Suspension Culture 90<1 (SUMITOMO) to make neurospheres (NSs). For suspension culture, we used NS medium containing media hormone mix (MHM) medium (KOHJIN-Bio) supplemented with 2% MACS NeuroBrew-21 (Miltenyi Biotec), 20 ng/ml basic FGF (PEPROTECH), 3 µM SB431542, 3 µM CHIR99021, and 10 ng/ml LIF (Millipore). The medium was changed every 3-4 days and NSs were passed every week. At passages, we added 3 µM Y-27632 (WAKO) to NS medium. After more than 3 passages, the dissociation step was performed.

Dissociated cells were plated onto mouse astrocytes (see above) at a density of 8 x 10^5^ cells per array. These cells were cultured in differentiation medium containing MHM supplemented with 2 % MACS NeuroBrew-21, 20 ng/ml BDNF (PEPROTECH), 10 ng/ml GDNF (PEPROTECH), 0.2 mM L-Ascorbic Acid (Sigma-Aldrich), 400 nM dibutyryl -cAMP (Sigma-Aldrich) and 2 nM DAPT (Sigma-Aldrich) for 28 days. Half the medium was changed twice a week.

### MEA recording with 2-D differentiated neurons

All MEA recoding was performed using MEA-2100 Systems with 60 electrode head stage (MEA2100-HS60) and Multi Channel Experimenter (Multi Channel Systems). All recordings from each electrode were acquired at 20 kHz and filtered at 200 Hz by a second-order high pass filter of Butterworth type. Observations of the spontaneous activities were performed for 5 min at room temperature on the head stage in a clean bench. Obtained spontaneous spike activities were analyzed by MC_Rack software (Multi Channel Systems). The threshold for spike detection was set to six standard deviations of the average signal-noise ratio. Burst activity was defined as follows: minimum (min) duration of a burst, 20 ms; min number of spikes in a burst, 4; maximum inter-spike interval in a burst, 10 ms; min interval between 2 bursts, 20 ms. Burst detection was performed individually for each electrode. Recordings were conducted until 35 days post-dissociation.

### Cortical organoid culture

For organoid culture, we referred to 3 protocols and combined them with some modifications (Trujillo *et al*, 2019; Pasca *et al*, 2015; Lancaster & Knoblich, 2014). Briefly, hESCs were cultured with DEF-CS medium under feeder-free condition. Then, these cells were dissociated using TrypLE select, and resuspended in STEP1 medium containing DEF-CS medium with 10 µM SB431542 and 1 µM dorsomorphin. About 4 x 10^6^ cells were transferred into 1 well of an Ultra-Low attachment 6-well plate (Corning) and kept in suspension under 90 rpm rotation. Only on day 0, the medium was supplemented with Y-27632, exchanged daily for 7 days and then substituted by STEP2 medium containing Neurobasal medium (Gibco) with 2 % MACS NeuroBrew-21 w/o Vitamin A (Miltenyi Biotec), 1X Glutamax (Gibco), 20 ng/ml Recombinant human FGF basic (PEPROTECH), 20 ng/ml EGF (Gibco), and 100 U/ml penicillin 100 µg/ml streptomycin (Nacalai). The medium was exchanged daily for 21 days. On day 28, each organoid of neural precursor cells (NPC organoid) was transferred to each Matrigel droplet (Corning) on a PARAFILM sheet (Bemis). After incubating at 37 °C for 30 min, these droplets were transferred at a density of 6 droplets per 1 well of a 6-well plate into STEP3 medium containing a 1:1 mixture of DMEM/F12 (Invitrogen) and Neurobasal with 0.5X N2 supplement (Invitrogen), 1X Glutamax, 0.5X NEAA (Gibco), 2.65 µg/ml insulin, 0.1 mM 2-mercaptoethanol (Sigma-Aldrich), 100 U/ml penicillin and 100 µg/ml streptomycin. STEP3 medium was supplemented with 1 % MACS NeuroBrew-21 without Vitamin A and cells were kept in suspension without rotation for 4 days. After that, STEP3 medium was supplemented with 1 % MACS NeuroBrew-21 and cells were kept under 80 rpm rotation. The medium was changed every 3 days.

### Measurement of NPC organoids

Culture protocol was described above (See Cortical organoid culture). Bright-field images of NPC organoids were obtained using a fluorescence microscope, BZ-X800 (Keyence) and circumference of each organoid was automatically measured using Keyence Hybrid Cell Count Module, BZ-H4C (Keyence). The thresholds for organoid detection were preset as follows: size, 6,000-100,000 µm^2^; circularity > 80%. Measurements were performed on day 2, 7, 17 and 27. For measurement of each cell size, fluorescent images by SOX2 were obtained and the major axis of each cell was automatically measured. The threshold for size detection was preset as follows: size > 3.4µm (−2.5SD). Measurements were performed on day 27. Statistical analysis was performed using Welch’s t-test followed by Bonferroni correction for multiple comparisons.

### RT-qPCR for NPC organoids

On day 25, NPC organoids were collected to extract RNA using TRI Reagent. To analyze differentiation stages, designed primers were synthesized by IDT with the following sequences:

qPCR_POU5F1_F: 5’ TTGGGCTCGAGAAGGATGTGGT 3’
qPCR_POU5F1_R: 5’ TGCATAGTCGCTGCTTGATCGC 3’
qPCR_PAX6_F: 5’ ACCACACCGGTTTCCTCCTTCACA 3’
qPCR_PAX6_R: 5’ TTGCCATGGTGAAGCTGGGCAT 3’
qPCR_EOMES_F: 5’ GGTTCCCACTGGATGAGACA 3’
qPCR_EOMES_R: 5’ TGCAGTCGGGGTTGGTATTT 3’
qPCR_NCAM1_F: 5’ GACCCCATTCCCTCCATC 3’
qPCR_NCAM1_R: 5’ TCTGGTCGAGTCCACGAAG 3’
qPCR_MAP2_F: 5’ GGATCAACGGAGAGCTGAC 3’
qPCR_MAP2_R: 5’ TCAGGACTGCTACAGCCTCA 3’

### Single cell RNA-Sequencing (scRNA-Seq) by NPC organoids

On day 27, NPC organoids were dissociated using TrypLE Select at 37 °C for 5 min. After passing through 20 µm strainers, cells were resuspended with STEP2 medium (see Cortical organoid culture) and kept at 4 °C. Cell concentration and viability were assessed using TC20 Automated Cell Counter (Bio-Rad).

Single cells were processed through the Chromium Single Cell 3’ Gel Bead, Chip and Library Kits v3.1 according to the manufacturer’s protocol (10X Genomics). Cells were added to each channel and immediately loaded and partitioned into 3’ Gel Beads in emulsion using Chromium Controller. After barcoded reversed transcription, gel beads in emulsions (GEMs) containing cDNA molecules were disrupted and purified and then PCR-amplified. After amplified cDNAs were fragmented, 5’ adapter and sample indices were incorporated as the final libraries. Concentrations of the amplified cDNA and size profiles of the final libraries were examined by Agilent Bioanalyzer 2100 using a High Sensitivity DNA chip (Agilent). Final libraries were quantified using KAPA Library Quantification Kits (Roche). The libraries were loaded at a concentration of 10 nM and sequenced on NovaSeq 6000 system (Illumina) with the following conditions: Paired-end, single indexing; Read 1, 28 cycles; i7 Index, 8 cycles; Read 2, 91 cycles.

Raw sequencing data were preprocessed with Cell Ranger software (version 3.1.0, 10X Genomics). After converting binary base call (BCL) files and demultiplexing via cellranger mkfastq pipeline, reads were aligned to hg19 human reference genome and the feature-barcode matrix was generated via cell ranger count pipeline. Information about key metrics such as number of total reads, estimated number of cells, mean reads per cell, median genes per cell, median unique molecular identifier (UMI) counts per cell and % of fraction reads in cells were also obtained via this pipeline.

Combining data of all 3 samples and normalizing the read depth before merging were performed via cell ranger aggr pipeline. Integrated datasets were obtained for data visualization and further analysis via Loupe Cell Browser software (version 4.0, 10X genomics). After removing clusters which had more than 50 % of mitochondrial genes (with ’MT-’ prefix) in TOP 20 expression genes, data were reanalyzed via cell ranger reanalyze pipeline. Uniform Manifold Approximation and Projection (UMAP) plots were displayed via Loupe Cell Browser software from this file and used to explore the integrated datasets. Validation of the optimal number of clusters in K-means clustering was performed via the clusGap function with cluster v2.1.0 package (Martin *et al*, 2019) after dimension reduction by principal component analysis (PCA) with stats v3.6.2 package (R Core team, 2020). A total of 10 clusters were generated, which were further merged into final 8 clusters based on the expression pattern of marker genes. Marker genes were mainly selected from Transcriptomic Explorer in Allen Human Brain Atlas and a published paper (Trujillo *et al*, 2019). Gene ontology (GO) analysis was performed using Metascape (Zhou *et al*, 2019) and ToppGene Suite (Chen *et al*, 2009a). Functional roles and related diseases of each gene were examined using GeneCards database (Stelzer *et al*, 2016).

### Immunohistochemistry

NPC organoids and cortical organoids were fixed with 4 % paraformaldehyde overnight at 4 °C and replaced to 10 %, 20 % and 30 % sucrose buffer. These were then embedded using O.C.T compound (Sakura) with 10 % sucrose. Embedded samples were sliced in a cryostat (10 µm slices). After air dry, these sliced samples were blocked and permeabilized with 0.3 % Triton-X-100 and 3 % normal goat serum (NGS) in PBS for 1 hour at room temperature. They were then incubated with primary antibodies diluted in solution containing 0.3 % Triton-X-100 and 3 % NGS overnight at 4°C. Following primary antibodies were used in this study: mouse anti-SOX2 (sc-365823, 1:200, SANTA CRUZ); rabbit anti-TBR2 (ab23345, 1:200, Abcam); mouse anti-B-Tubulin (MMS-435P, 1:200, COVANCE); rabbit anti-CTIP2 (ab28448, 1:1000, Abcam). Then these sliced samples were washed with 0.05 % Tween 20 (Nacalai) 3 times and incubated with secondary antibodies (Alexa Fluor 488- and 561-conjugated antibodies, Life Technologies, 1:500) diluted with 0.05 % Tween 20 for 1 hour at room temperature. Nuclei were visualized with DAPI solution (0.1 µg/ml). These sections were mounted using VECTASHIELD Antifade Mounting Medium (Vector) and analyzed under a confocal microscopy (FV3000 scanning confocal microscope, Olympus).

### Statistical Analysis

Sample size was determined based on the experimental designs except for measurement of NPC organoid. As for measurement of NPC organoid, effect size and standard deviation were estimated according to the pilot study and the sample size was calculated by T statistic using Sample Size Calculators (UCSF CTSI, https://sample-size.net/). Each data was obtained from different samples. For statistical analysis, Welch’s t-test with post hoc Bonferroni correction, one-way ANOVA with post hoc Tukey-Kramer test and Mann-Whitney U test with post hoc Bonferroni correction were performed using stats v3.6.2 package (R Core team, 2020).

## Author contributions

Conceptualization: Y.N., J.N. and T.T., Methodology: Y.N. and J.N., Validation: Y.N. and J.N., Investigation: Y.N., J.N. and T.N., Supervision: T.T., Writing - Original Draft: Y.N., J.N., T.N. and T.T., Funding Acquisition: T.T.

## Acknowledgements

We thank Yuriko Kusakari, Kyoko Shiga and all technical staffs in Takumi lab for technical assistance. We thank Kota Tamada for helpful advises in using the confocal microscope and the R software. We also thank Manabu Toyoshima and Yasuko Hisano for helpful advises and demonstrations about transfection and cell differentiation procedures. RT-qPCR was performed using QuantStudio 12K Flex, Common Use Equipment, in the Support Unit for Bio-Material Analysis, Research Resources Division, RIKEN Center for Brain Science. This work was supported by RIKEN Junior Research Associate Program. This work was supported in part by KAKENHI (16H06316, 16H06463) from Japan Society for the Promotion of Science and Ministry of Education, Culture, Sports, Science, and Technology, Intramural Research Grant for Neurological and Psychiatric Disorders of NCNP, the Takeda Science Foundation and Smoking Research Foundation.

## Data availability

The scRNA-seq data reported in this paper are available in the DDBJ/EMBL/GenBank databases (under submission). All supplementary information is available in the online version of this article. Further information and requests for materials should be addressed to the corresponding author.

## Supplementary Figure legends

**Supplementary Figure 1. aCGH assays of whole chromosomes for (A) 1q del and (B) 1q dup clones.**

Gains or losses were determined by normalized log10 ratios. Although there were some plots distant from the base line besides 1q21.1 region (such as red dots on chromosome 16), the number of plots within these regions were few (<= 5), which meant very short ranges. These plots were mostly common between 2 genotypes.

**Supplementary Figure 2. Relative RNA expressions of targeted and neighboring genes in (A) 1q del and (B) 1q dup clones.**

Reduction or increase of gene expressions were observed within the range of target regions in (A) 1q del and (B) 1q dup, respectively. Data are normalized by CTRL scores and shown as mean ± SEM (n=2 (HJV, NUDT4B), n=3 (PRKAB2, BCL9, GPR89B)); ***p<0.001; one-tailed Welch’s t-test.

**Supplementary Figure 3. MEA analysis for spike activities overtime and bursts.**

(A) Median spike frequency in top 20 electrodes of a 60-6well MEA dish over time. Data are shown as bee swarm plots (n = 20 each). Black open circles represent electrodes with no spikes. Black horizontal lines show median scores. (B) Median burst frequencies of each genotype at 35 days post-dissociation showing significantly more bursts in 1q del than CTRL. Top 5 data from 2 batches (circle, 1-well array; square, 6-well array) of each genotype were extracted. Data are shown as bee swarm plots (n=10 each). Black horizontal lines show median scores; **p<0.01; Mann-Whitney U test with Bonferroni correction as post-hoc analysis.

**Supplementary Figure 4. Alkaline phosphatase staining to check pluripotency.**

(A) A macroscopic image of stained hESCs cultured on a plastic well. Wild-type cells before transfection. Scale bar, 5 mm. (B) Bright field images of stained hESCs. 3 cell lines (CTRL, 1q del and 1q dup) after targeting procedures. Scale bar, 100 µm.

## Supplementary Table legends

**Supplementary Table 1. Enrichment Analyses with Disease Ontologies.**

**Supplementary Table 2. Functions and related diseases of each gene in Common DEGs.**

**Supplementary Table 3. Pathway analyses by NSC cluster.**

